# Reproductive effort of intertidal tropical seagrass as an indicator of habitat disturbance

**DOI:** 10.1101/2020.07.19.177899

**Authors:** Amrit Kumar Mishra, Deepak Apte

**Affiliations:** Marine Conservation Department, Bombay Natural History Society, Hornbill House, Dr. Salim Ali Chowk, Shaheed Bhagat Singh Road, Opp. Lion Gate, Mumbai, 400001, India

**Keywords:** Turtle grass, *Thalassia hemprichii*, habitat disturbances, ecological indicator, reproductive ecology

## Abstract

Habitat disturbance is one of the major causes of seagrass loss around the Andaman Sea in the Indian Ocean bioregion. Assessing the seagrass response to these disturbances is of utmost importance in planning effective conservation measures. Here we report about seagrass reproductive effort (RE) as an indicator to assess seagrass response to habitat disturbances. Quadrat sampling was used to collect seagrass samples at three locations (a disturbed and undisturbed site per location) of the Andaman and Nicobar Islands. The health of seagrass meadows was quantified based on density-biomass indices of the disturbed sites. A change ratio (D/U) was derived by contrasting the RE of disturbed (D) sites with the undisturbed (U) sites of all three locations. The relationship between RE and plant morphometrics were also quantified. Reproductive density of T. *hemprichii* was higher and significant at the three disturbed sites. The average reproductive density of *T. hemprichii* at the disturbed sites was 3.3-fold higher than the undisturbed sites. The reproductive density consisted around 52% of the total shoot density of *T. hemprichii* at the disturbed sites. In general, the increase in the plant RE was site-specific and was 4-fold higher at the three disturbed sites. Positive and significant correlations was observed between the change ratio of RE and the plant morphometrics, suggesting an active participation of seagrass morphometrics in the reproductive process. Increase in seagrass RE can contribute to the increase in population genetic diversity, meadow maintenance and various ecosystem functions under the influence of anthropogenic disturbance scenarios.

## 1. Introduction

Globally seagrass ecosystems are under various stress due to accelerating degradation of their habitats as a consequence of both direct and indirect human activities and climate change (Waycott et al., 2009; Short et al., 2011; Mckenzie et al., 2020). Natural events such as cyclones, hurricanes, tsunamis (Carlson et al., 2010; Côtè-Laurin et al., 2017) and diseases contribute to the decline of seagrass ecosystems to some extent, but majority of the seagrass loss are related to anthropogenic disturbances (Waycott et al., 2009; Short et al., 2011; Unsworth et al., 2019). These anthropogenic disturbances include coastal constructions, trawling, dredging, waste water disposal and aquaculture expansion that have led to loss of seagrass habitats (Kaldy and Dunton, 2000; Erftemeijer and Lewis, 2006; Short et al., 2011). Globally around 30% of seagrass population has been under decline due to habitat loss in the last three-four decades, with a yearly average loss of around 7% (Waycott et al., 2009; Pendleton et al. 2012; Howard et al. 2014). However, much of this decline has occurred in the Indian Ocean bioregion hotspot of seagrasses in the Andaman Sea, Java Sea, South China Sea and Gulf of Thailand (Short et al. 2011; Saenger et al. 2012). These scenarios call for an increase in understanding of the plant response to various local habitat disturbances, to plan efficient management and conservation measures.

Seagrass sexual reproductive effort (RE) is considered as an ecological indicator of various natural and anthropogenic disturbances (Collier and Waycott, 2009; Cabaço and Santos, 2012; Ali et al., 2018; Lawrence and Gladish, 2019).Through increased RE the plant can recover from the loss occurred due various disturbances induced stress (Collier and Waycott, 2009; Cabaço and Santos, 2012; McKenzie et al., 2016) by allocating higher plant resources for production of flowers/fruits (Kaldy and Dunton, 2000; Smith et al., 2016) that invariably helps the seagrass to maintain the meadow genetic diversity (Ackerman, 2006; McMahon et al., 2017). Recovery of seagrass meadows from these disturbances are directly dependent on the increased RE of the plant to produce seed banks (Hammerstorm et al., 2006; Cabaço et al., 2009; McKenzie et al., 2016). Increased RE can also enhances the resilience and probability of adaptation of the seagrass meadows to changing climatic conditions (Ehlers et al., 2008).

Thalassia (Hydrocharitaceae) is a widely distributed seagrass in both tropic and temperate regions of the world having two species; *Thalassia testudinum* and *Thalassia hemprichii* (Short et al., 2011; McKenzie et al., 2020). *T. hemprichii* is one of the keystone seagrass species that is dominant around the Indian Ocean bioregion including the waters of Andaman Sea (Short et al., 2011; Saenger et al. 2012). *T. hemprichii* is one of the persistent species, relatively slow-growing and resistant to anthropogenic pollution (Kilminster et al., 2015). However, when it comes to habitat disturbance, its vulnerability increases due to loss of plant morphometric features and population longevity (Mishra and Apte, 2020). Secondly, when *T. hemprichii* is present in shallow intertidal regions (<1m), the influence of habitat disturbances is severe on the plant vegetative propagations (Agawin et al., 2001; McMahon et al., 2017; Mishra and Apte, 2020), that increases the significance of reproductive processes for meadow maintenance.

Various types of habitat disturbances and their influence on the RE of Thalassia species have suggested that plant response to both natural and anthropogenic habitat disturbances include a positive response through increase in sexual reproduction (Cabaço and Santos, 2012; Ali et al., 2018; Lawrence and Gladish, 2019). The natural disturbances such as hurricanes (Gallegos et al., 1992; van Tussenbroek, 1994), heavy rainfall and terrestrial run-off (Chollett et al., 2007; McDonald et al., 2016), dredging (Kaldy and Dunton, 2000) and intertidal, nutrient and temperature stress (Durako and Moffler, 1985; Philips et al., 1981; Kelly and Dunton, 2017) have indicated a positive response on *T. testudinum* flower/fruit production or increase in reproductive shoots. However, most of the studies on *T. testudinum* are geographically confined to the Florida and Texas coast of the USA (Phillips et al., 1981; Duarko and Moffler, 1985; Kaldy and Dunton, 2000; Whitefield et al., 2004; McDonald et al., 2017; Kelly and Dunton, 2017), the Mexican Caribbean (Gallegos et al., 1992; van Tussenbroek, 1994) and Venezuela (Chollett et al., 2007).

Consequently, the effects of various habitat disturbances on RE of *T. hemprichii* are concentrated along the coast of Indonesia and Philippines surrounding the bioregion around the Andaman Sea (Agawin et al. 2001; Rollon et al. 2001; Olesen et al., 2014; McMahon et al., 2017). The various types of habitat disturbances on *T. hemprichii* include cyclones and grazing (Olesen et al., 2014; McMahon et al., 2017), land based run-off combined with environmental stressors (Rollon et al., 2001; Lwarance and Gladish, 2019) and mechanical damage (Olesen et al., 2014) that exerted a positive effect on the RE of the plant. However, there is not a single report on *T. hemprichii* reproductive effort from the coast of India including the Andaman and Nicobar Islands of the Andaman Sea, even though *T. hemprichii* is one of the keystone species found in the shallow intertidal regions of the Indian coast (Thangaradjou and Bhatt, 2018; Mishra and Apte, 2020).

In India, seagrass beds are declining due to various habitat disturbances (Nobi et al. 2011; Thangaradjou and Bhatt, 2018; Kaladharan and Anasukoya, 2019; Mishra and Apte, 2020) and in this scenario lack of a new methods to assess the effects of these disturbances on seagrass population can lead to severe loss of these ecosystems. Other than habitat disturbances due to human induced activities, natural phenomenon such as frequent cyclones (of the East coast of India) and monsoon rains (from West coast of India) and associated land-run-offs also affect the seagrass population structure.

13 out of 16 seagrass species found in India are recorded from ANI (Thangaradjou and Bhatt, 2018; Ragavan et al., 2016) and *T. hemprichii* is widely distributed species around Andaman Sea (Short et al., 2011; Ragavan et al., 2016; Mishra and Mohanraju, 2018). However, it is highly susceptible to long term habitat disturbances and its recovery periods are usually longer due to its slow growth patterns (Kilminster et al., 2015; Mishra and Apte, 2020). Loss of *T. hemprichii* meadows will deprive the sourrounding ecosystems of various important ecosystem functions, such as loss of feeding habitats for the endangered sea cows (*Dugong dugon*) and sea turtles (Murugan, 2004), loss of nurseries for commercially important fish population (Unsworth et al., 2018) and loss of carbon sequestration and storage in the coastal ecosystems (Howard et al., 2014; Dahl et al., 2016) and subsequent negative impacts on the coastal human communities of livelihood options.

Therefore, the present study will quantify the reproductive ecology *T. hemprichii* from three locations of the Andaman and Nicobar Islands (ANI), that are under the influence of habitat disturbance. We will derive the health status of these disturbed seagrass meadows using the widely accepted density-biomass relationship index. The reproductive density, total shoot density and morphological features of the plant will be measured. The change in RE derived from disturbed sites will be contrasted with undisturbed sites at each study location to assess i) the general RE trend of *T. hemprichii* to various disturbances across the three locations, ii) variation in RE with respect to locations and iii) if RE correlates with plant size and growth. We will also assess if change in RE of *T. hemprichii* of ANI can be used as an indicator of habitat disturbances.

## 2. Materials and Methods

### 2.1 Study sites

We surveyed the *T. hemprichii* meadows of Neil (now officially called Shahid Dweep), Havelock (now officially called Swaraj Dweep) Islands and Burmanallah location of ANI Hereafter we will use Neil, Havelock Islands and Burmanallah for convenience in the entire text. We surveyed the Neil, Havelock Islands and Burmanallah location of ANI of India during the months Feb-March 2019 when there is peak in flower/fruit production (Jagtap et al., 2003). These study locations exhibited a semi-diurnal tidal amplitude of 2.45m. The climate of these islands is typically tropical, with heavy rainfall, cyclones and hot and humid conditions (Sahu et al., 2013). However, we sampled in the dry season, when there was no land run-off except at the Burmanallah location.

#### 2.1.1. Neil Island

Neil Island is situated in the south-east region of ANI (Fig.1) The *T. hemprichii* population at the disturbed site (site 1) was under the influence trampling and tourism related disturbances. The undisturbed site (site 2) was 1000m apart from site 1 separated by dead coral (back reef) patches. The site 2 was under sheltered conditions due to the presence of dead coral pools. *Thalassia hemprichii* population was sampled at both sites within a depth of 0.5 m during low tide.

**Fig. 1.**
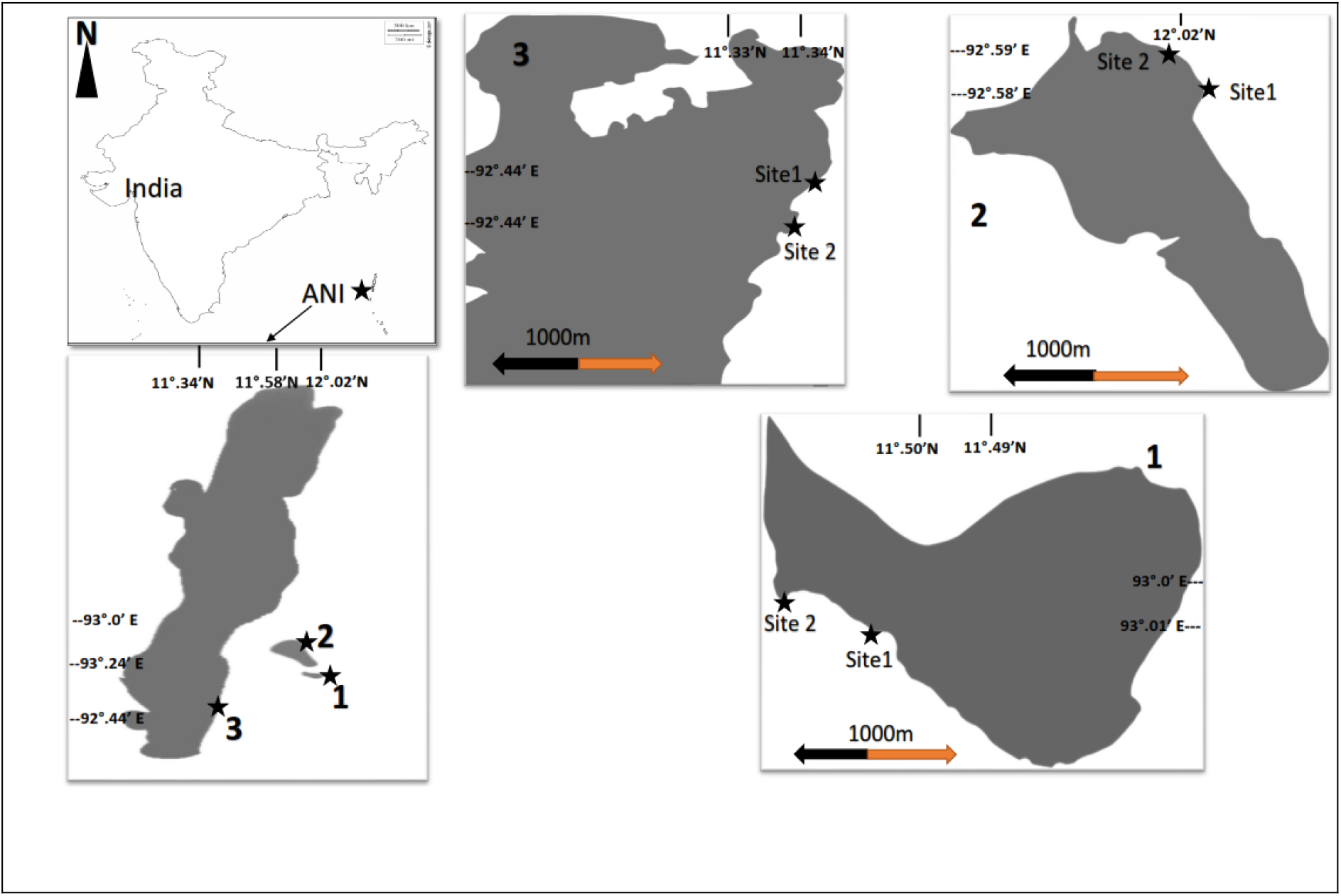
Map showing the study locations of Neil (1), Havelock islands and Burmanallah (3) of ANI, India. Each location has a disturbed (site 1) and undisturbed (site 2) study sites.

#### 2.1.2. Havelock island

Havelock Island is also situated in the south-east of ANI and north of Neil Island (Fig.1). The *T. hemprichii* population at site 1 was under the influence of physical damage due to boat anchoring and trampling due to tourism activities. The site 2 was 1000m apart and was surrounded by dead coral (back reef) patches. *T. hemprichii* population was sampled within a depth of 0.5 m during low tide.

#### 2.1.3. Burmanallah

Burmanallah is situated in the south-east region of ANI (Fig.1). This location has rocky intertidal beaches and human-made coastal concrete walls. The site 1 was under the influence of boat anchoring and trampling damage along with coastal developments, agricultural land run-off and domestic waste water discharge from the nearby village. The site 2 was 1000m apart from site 1 and sheltered by rock pools. *T. hemprichii* population was sampled within a depth of 0.3m during low tide.

### 2.2. Water and sediment sampling and analysis

Physical parameters like pH, temperature and salinity were measured in the field using portable sensors from the same site where sediment and seagrass quadrats were collected. Temperature and pH were measured using a portable pH meter (HI991300P, Hanna Instruments, Germany) and salinity with salinometer (HI98203, Salintest, Hanna Instruments, Germany). Sediment cores (n=10) were collected from each quadrat of all six sites where seagrass was sampled, using a 5 cm wide and 10 cm long plastic corer. The corer was pushed at least 5 cm into the sediment while sampling. As the sampling sites were full of coral rubbles, it was difficult to dig deeper. Sediments were collected in plastic bags and transferred to the laboratory. In the laboratory, the sediment samples were oven dried at 60°C for 72 hours before sieving for various grain sized fractions (500, 150, 75, 63μm).

### 2.3. Seagrass sampling and analysis

Quadrats (n=10) were collected randomly from a transect of 20 × 30m^2^ perpendicular to the beach at both the sites (disturbed and undisturbed) from all three locations, within a depth of 0.3 to 0.5m during low tide. We used a quadrat of 30cm^2^ and a hand shovel to dig out seagrass samples up to 10 cm depth. From each quadrat seagrass collected were rinsed with seawater carefully (without dislocating the rhizome mat) in the field and brought to the lab for further analysis. In the laboratory samples were washed gain with fresh water and epiphytes were removed with a plastic razor. In each sample, the total density (individuals m^−2^) was estimated by counting the number of shoots of physically independent individuals. The presence of male and female reproductive structures like flowers or fruits, in each shoot was recorded to estimate the reproductive shoot density (fruits/flowers m^−2^). Morphometric variables such as maximum leaf length (n=15/quadrat/site), rhizome diameter (n=20/quadrat/site) was measured using a Vernier Calliper (accuracy:0.02mm) and a meter tape. The leaves, roots, vertical and horizontal rhizomes were separated and oven dried for 48 h at 60° C for total biomass and shoot biomass (vertical + horizontal biomass) estimates (g DW m^−2^). Seagrass abundance (density-biomass relationship) was used to quantify the health status of the *T. hemprichii* meadows. We used these indices as they are used globally (Marba et al., 2013; Vieira et al., 2018) and locally around the Andaman Sea (Ali et al., 2018).

Horizontal rhizome elongation rate was estimated for each site based on the linear regression of the number of horizontal internodes in between two consecutive shoots against the age difference of these shoots (Mishra and Apte, 2020). The slope (internodes yr^−1^) was then multiplied by the mean internodal length (cm internode^−1^) to obtain the horizontal elongation (cm yr^−1^). For each site only one horizontal elongation value was obtained, as other than quadrat samples, few plant rhizomes were hand-picked for better representation of horizontal elongation. Reconstruction techniques an indirect measurement of plant growth (Duarte et al., 1994; Fourqurean et al., 2003) was used to derive plastochrome intervals required to estimate the shoot age of the plants (Mishra and Apte, 2020).

The reproductive effort (RE %) was calculated by dividing the reproductive density by the total shoot density multiplied by 100. A change ratio, D/U, was used to quantify the RE response to various disturbances (Cabaço and Santos, 2012), where D is the RE of the disturbed site (site 1) and U is the RE of the undisturbed site (site 2). The change ratio indicates the relative increase in RE for the disturbed seagrass plants. The changes in RE were classified into two orders, i) increasing or decreasing, if the RE changed by >5%, and ii) no change, if the RE changed ≤5%.

### 2.4. Statistical analysis

For density-biomass relationship, log10 of both density and biomass was plotted against each other using linear regression. A higher regression coefficient value (r >0.80) indicates healthy meadows and a lower value (r<0.50) indicates unhealthy meadows (Vieira et al., 2018).

A Chi-square test (p<0.05) was used to test the deviation from each partitioning of the trends of RE. The effects of both size (leaf length and rhizome diameter) and growth (biomass and horizontal elongation rate) of *T. hemprichii* on the change ratios of RE was tested using linear regression analysis (Sokal and Rohlf, 1995).

Significant differences between *T. hemprichii traits* reproductive density and RE estimates were investigated using two-way ANOVA with sites (two sites) and locations (three locations) as fixed factors. All data was pre-checked for normality (Shaphiro-Wilks test) and homogeneity of variance (F-test). When variances were not homogenous, the corresponding data was ln (x+1) transformed. When there were significant interaction effects, the Holm-Sidak test was performed for a posteriori comparison among factor levels (sites and locations). All statistical analysis was carried out using SIGMAPLOT (Ver. 11.02) software.

## 3. Results

The disturbed sites had relatively low pH levels compared to their respective undisturbed sites. The salinity levels between the sites and locations did not show much variation. In general, the sediment silt content was significantly higher at the disturbed sites than the undisturbed sites. The highest silt content (27.86±0.07%) was observed at disturbed site of Neil Island and the lowest (14.73±2.13%) at the disturbed site of Havelock Island (Table 1).

The density-biomass relationship indicated that habitat disturbance has a negative effect (unhealthy)on the *T. hemprichii* meadows at the site1 of each three locations. This was evident from the low correlation coefficient value (R^2^=0.31, p<0.001) obtained from the linear regression of density-biomass relationship (Fig.2).

**Fig.2.**
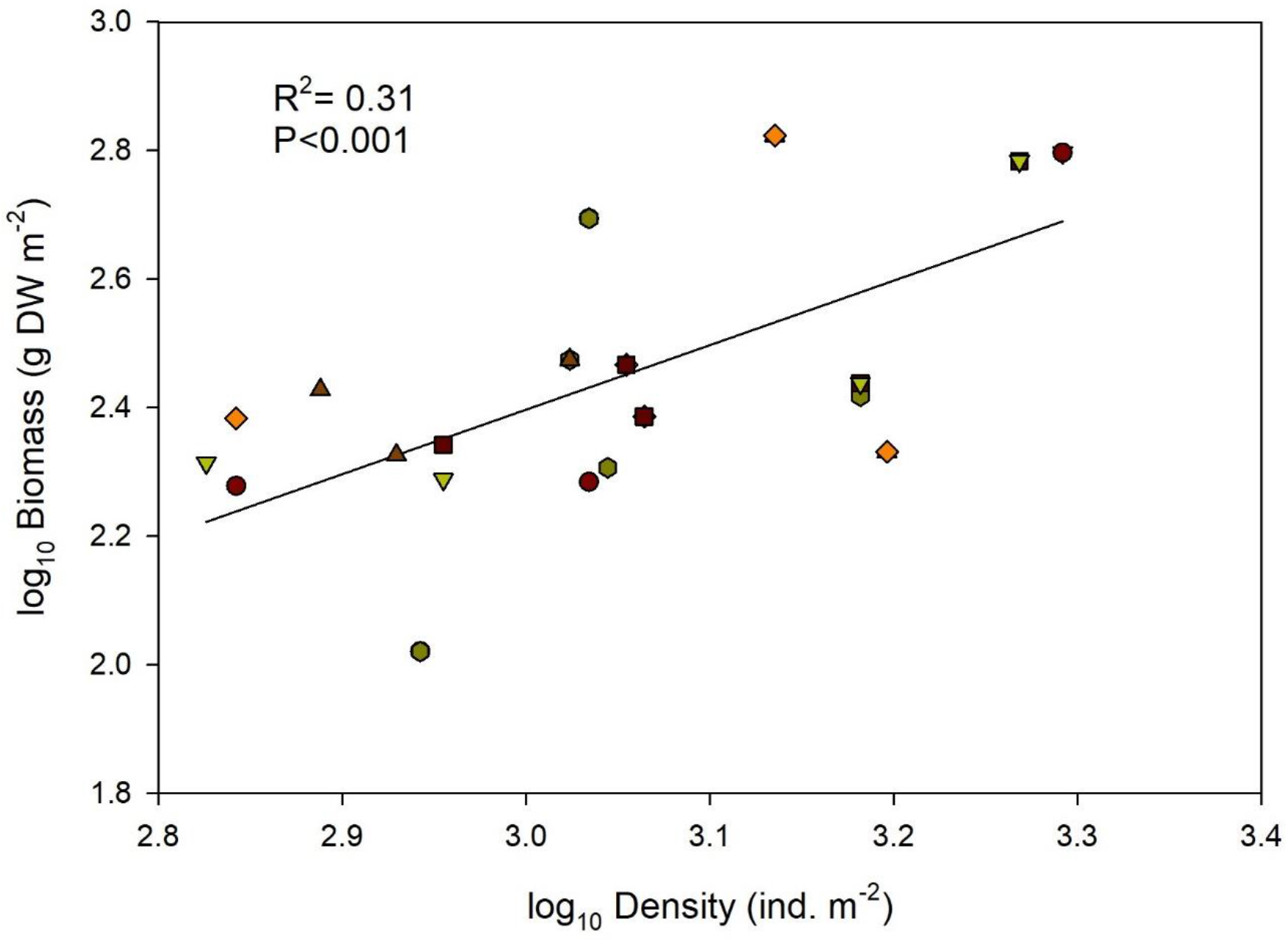
Density-Biomass relationship of *T. hemprichii* meadows at the disturbed sites of Neil (●) Havelock (▼) islands and Burmanallah (■) of ANI. All quadrats (n=30/3 sites) were pooled together to have a better representation

Reproductive density of T*. hemprichii* was higher and significant at the disturbed sites of all three locations (Fig. 2). The reproductive density of *T. hemprichii* at the Burmanallah disturbed site (site 2) was 4-fold higher than the undisturbed site, which was the highest among the three locations (Fig. 2A). The reproductive density of *T. hemprichii* at the disturbed sites of the Havelock Island consisted 64% of the total shoot density, whereas the disturbed sites of the Burmanallah and the Neil Island accounted for 50% and 41% of the total shoot density respectively.

A magnitude of positive response was observed for T. *hemprichii* RE with various plant morphometric and growth features. *T. hemprichii* responded positively to habitat disturbances through increase in RE (Fig. 2B). This increase in RE was higher for the disturbed site of the Havelock Island and the Burmanallah than that of the Neil Island. There was a 4.5-fold (the highest among the locations) increase in RE at the disturbed site of the Havelock Island, followed by a 4.4-fold increase at the Burmanallah and 3.5-fold increase at the disturbed site of Neil Island (the lowest among the locations). The change ratio (D/U) followed a similar pattern of increase related to RE. The change ratio increased 6-fold at the disturbed site of the Havelock Island, followed by a 5.5-fold increase at the Burmanallah and 4-fold increase at the Neil Island (Fig.2B).

Under the influence of disturbed conditions, a common pattern of increase in seagrass morphometric (growth and size) features was observed with increase in RE of the plant under disturbed conditions. The horizontal elongation rate was strongly (R^2^=0.98) and significantly (P=0.026) correlated with the plant RE at all three disturbed sites (Fig.3). The highest horizontal elongation rate was observed at the Neil Island (with the lowest D/U) and the lowest elongation rate at the Havelock Island (with the highest D/U) (Fig.4a). A significant positive response of change in RE to the rhizome diameter of the seagrass was observed at the Neil, Havelock Islands and the Burmanallah disturbed sites (Fig.4b). The highest increase in rhizome diameter of *T. hemprichii* in relation to the change ratio of RE was observed at the Neil Island, whereas similar levels of increase in rhizome diameter was observed at the Havelock Island and the Burmanallah in relation to the change ratio of RE (Fig.4b).

**Fig.3.**
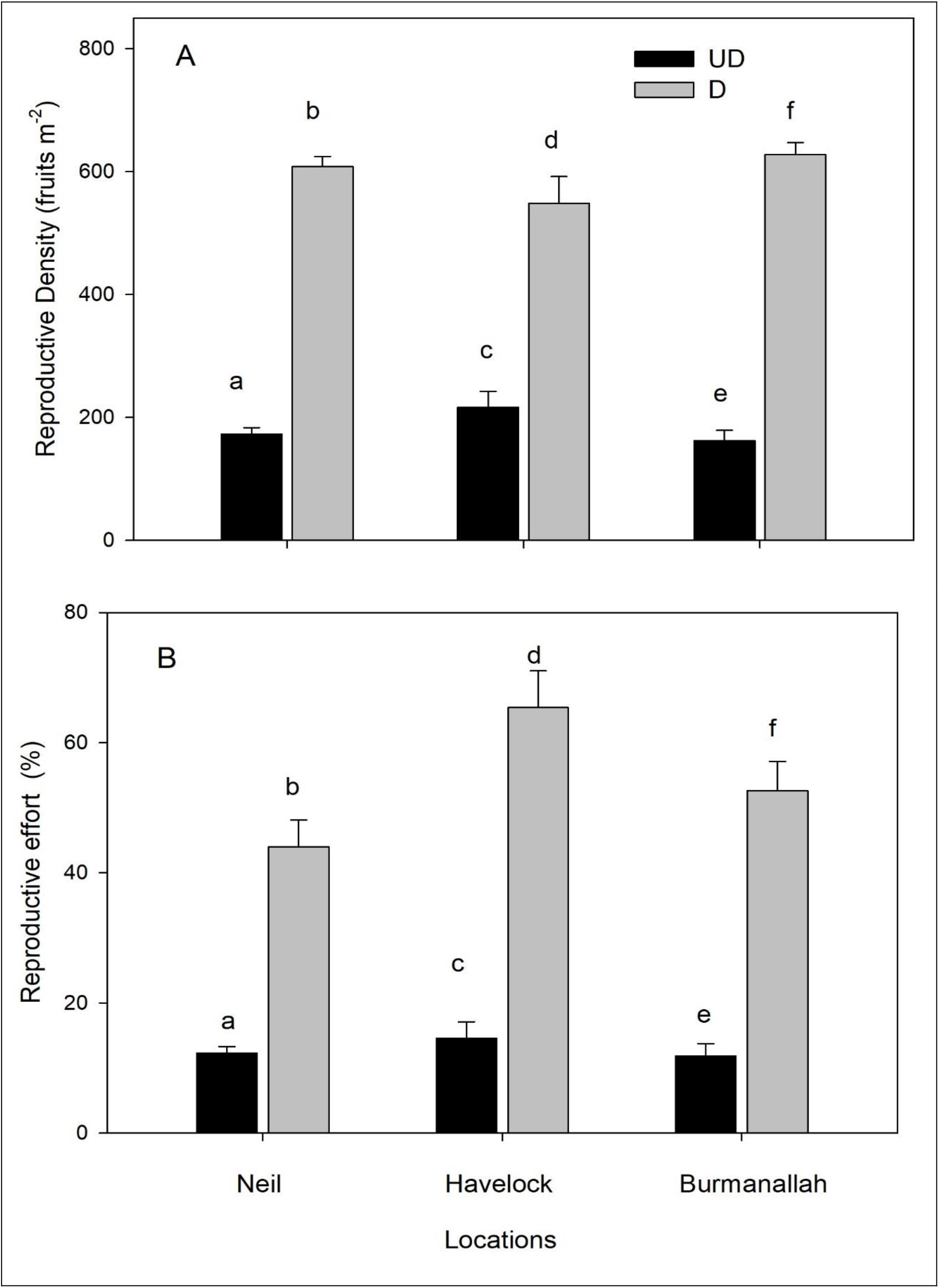
A) Reproductive density and B) reproductive effort (%) of *T. hemprichii* under disturbed (D) and undisturbed (UD) sites at the three locations of ANI. Error bars represent standard errors. Small letters indicate significant differences between sites (D and UD).

**Fig. 4.**
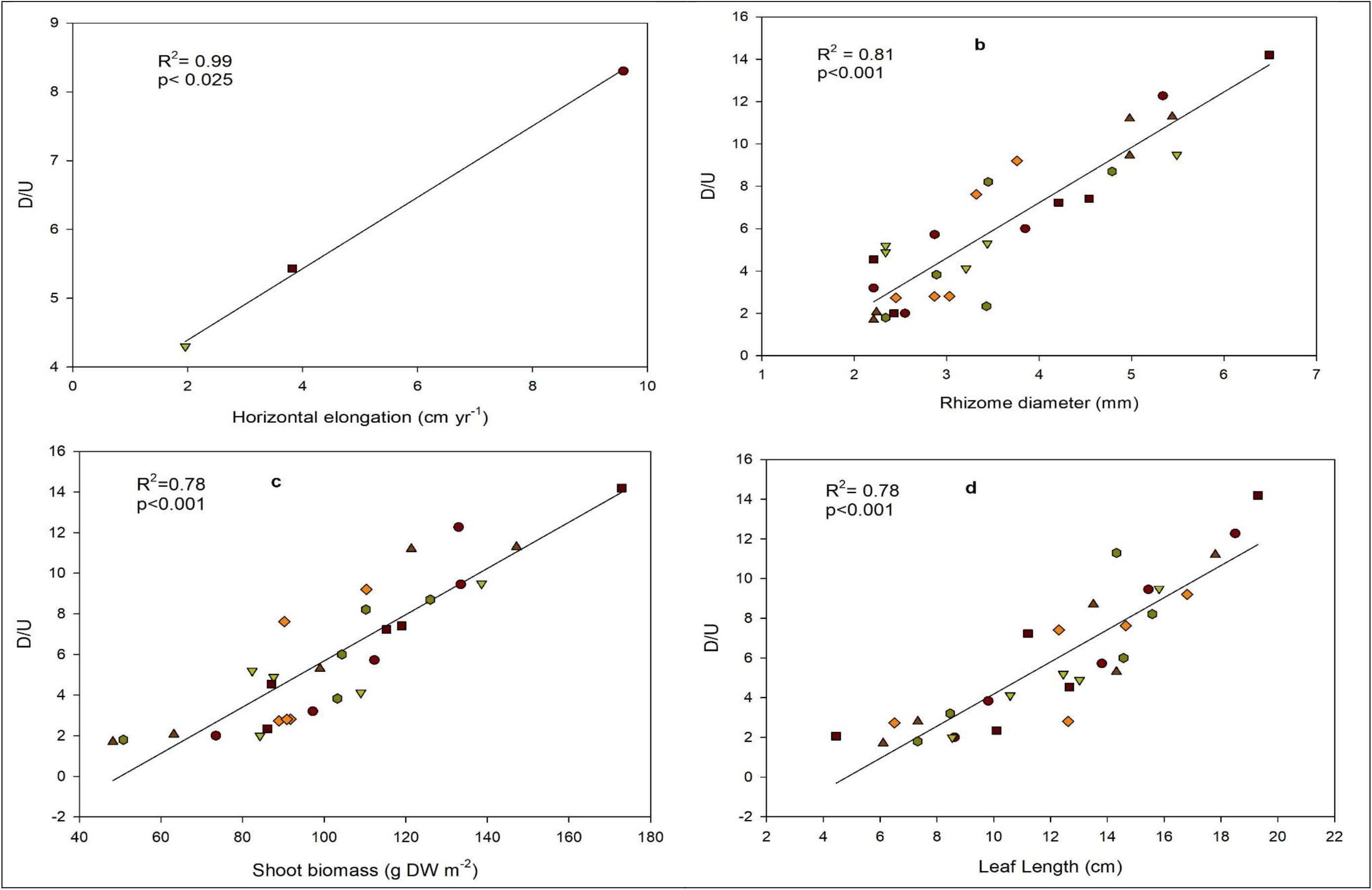
Relationship between mean change ratio (D/U) of the reproductive effort of *T. hemprichii* and a) horizontal elongation, b) rhizome diameter, c) shoot biomass and d) maximum leaf length responding positively to various disturbances at Neil (●) Havelock (▼) islands and Burmanallah (■) of ANI. Except a) horizontal elongation, for b, c and d all quadrats (n=30/3 sites) were pooled together for better representation.

*T. hemprichii* shoots biomass and maximum leaf length also responded positively to the change ratio of RE at the disturbed sites (Fig.4c & d). The highest increase in shoot biomass and maximum leaf length with response to change ratio of RE was observed at the Neil Island, whereas the lowest shoot biomass and leaf lengths was observed at the disturbed site of the Burmanallah and the Havelock Island respectively (Fig.4c & d).

## 4. Discussion

Disturbance is an important driver of increasing tropical seagrass richness and seagrass respond to both natural and anthropogenic disturbance/stress by increasing their RE (Cabaço and Santos, 2012; McMahon et al., 2016) and we observed a positive response of seagrass *T. hemprichii* RE under the influence of habitat disturbances. Increase in RE is a general plant response observed for many terrestrial (Pascarella, 1998) and aquatic plants (Trèmolierès, 2004) this plant response may contribute to the increase in population genetic diversity (Bell et al., 2008; Lawrence and Gladish, 2019). Consequently, for the maintenance of the genetic diversity, the seeds developed during the reproductive process needs to be successful in i) bottom dispersal from the disturbed sites (Lacap et al., 2002) and ii) establishing new individual plants under the disturbed conditions (Karlson and Mendèz, 2005). Previous studies have observed this phenomenon of *T. hemprichii*, being successful in the establishment of new meadows from dispersed seeds along the coast of Philippines in the Andaman Sea (Rollon et al., 2001; Lacap et al., 2002). Being successful in establishing new plants/meadows may also enhance the plant resilience to various changes occurring in future and maintaining various ecosystem functions (Ehlers et al., 2008; Hughes and Stachowicz, 2009).

This study has highlighted two key stressors combined with habitat disturbance that affect seagrass RE; such as hydrodynamics and anthropogenic pollution. Anthropogenic pollution is increasing due to human induced activities such as coastal developments, increased tourism and waste water disposal (Waycott et al., 2009; Unsworth et al., 2017; Mishra and Kumar, 2020; Mishra et al., 2020 under review). However, change in hydrodynamics can be related to climate change; frequent weather changes that can intensify cyclones or hurricanes and associated wave dynamics (Sobel et al., 2016). Anthropogenic pollution is one of the major contributors to seagrass decline worldwide (Waycott et al., 2009; Short et al., 2011; Saenger et al. 2012) and in India (Nobi et al. 2010; Kaladharan and Anasukoya, 2019). Interestingly we observed that, the RE of *T. hemprichii* increased (ca.4.13-fold) under the influence of habitat disturbance combined with anthropogenic disturbances. This increase in RE of *T. hemprichii*, observed under the influence of physical damage (due to boat anchors and trampling) coincides with the response of *T. hemprichii* from the coast of the Philippines (Olesen et al., 2004), Indonesia and Australia (McMahon et al., 2017). Similar effects of habitat disturbances have been observed for and for *T. testudinum* along the coastal of the USA (Kaldy and Dunton, 2000; Whitefield et al., 2004) and the Mexico (Gallegos et al., 1992; van Tussenbroek, 1994).

The increase in RE of the seagrass under the influence of habitat disturbance combined with anthropogenic stressors (at the Burmanallah disturbed site) also coincides with previously observed results of *T. hemprichii* from the coast of the Philippines (Rollon et al., 2001) and Australia (Lwarance and Gladish, 2019). Similar results for *T. testudinum* have been observed from the coast of the USA under the influence land run-off (McDonald et al., 2016). Other than T. hemprichii and T. testudinum, various other seagrass species has also shown similar response to habitat disturbances and anthropogenic stressors through increase in their RE (Cabaço and Santos et al., 2012).

Physical damage by boat anchors and the effect of wave hydrodynamics causes loss of seagrass leaves, rhizome structures and breakage of flowers and fruits that can result in low reproductive density at the disturbed sites (Unsworth et al., 2017; Mishra and Apte., 2020). Thus, an increase in RE can be an adaptative response of *T. hemprichii* to these physical damages, which has been observed as a common response for various other seagrass species (Ehlers et al. 2008; Cabaço & Santos, 2012; Pereda-Briones et al. 2018). Being present in shallow coastal waters (<0.5m) *T. hemprichii* is also sensitive to burial stress due to sand wave hydrodynamics. To overcome this stress (combined with habitat disturbance and land run-off) *T. hemprichii* undergoes morphological changes by increasing leaf and rhizome lengths (Duarte et al., 1997; Mishra and Apte, 2020). Thus, this explains the positive relation between increase in RE and higher horizontal elongation rate of the plant in an attempt to migrate away from the disturbed conditions. Consequently, this migration of the plant will also depend on other abiotic factors such as nutrient limitation and sediment characteristics of the sites (Agawin et al., 2001; McMahon et al. 2017; Ali et al., 2018; Mishra and Apte, 2020). *T. hemprichii* prefers sandy habitat (Jagtap et al., 2003) and the disturbed sites with higher percentage of silt can also restrict this migration, as a result the plant invests in vertical leaf growth, which explains the positive relationship of RE with leaf length.

Intertidal stress due to high temperature (35-37°C) can be a major factor in reduction of *T. hemprichii* flower/fruit production (McMillian, 1980; Agawin et al., 2001; Pederson et al., 2016) which can lead to an increase the RE of the plant. The effect of high temperature on *T. hemprichii* have been reported from the coast of Kenya (McMillan, 1980) and for *T. testudinum* from the coast of the USA (Philips et al., 1981; Duarko and Moffler, 1985; McDonald et al., 2016). Importantly, in both species an increase in flowering/fruit production or maturation of fruits for release have been observed with increase in temperature. However, this response of increase in RE is species-specific (Cabaço and Santos, 2012) and plants with thinner rhizomes such as *Zostera noltii* (Cabaço et al., 2009) have shown a decrease in RE, whereas *Cymodocea racemose* and *Halodule uninervis* (Rasheed et al., 2004) have shown no response to temperature stress and *Halophila ovalis* and *Syringodium isoetifolium* (Rasheed et al., 2004).The disturbed sites (site 1) in our study were more prone to the intertidal temperature compared to the sheltered undisturbed sites (as they have higher water levels being present in rock pools or dead coral pools) during the daytime receding tides. This small temperature difference can result in reduced net photosynthetic production and increased photorespiration (Pedersen et al., 2016; Brodersen et al., 2019), which can deplete the plant energy sources required to increase their reproductive density. However, the below ground shoot biomass can contribute to the plant reproductive processes during this stress period as it stores higher energy (thus the positive correlation with RE at all three locations) and the roots can supply oxygen during daytime by passive diffusion from sediment (Borum et al., 2006) to maintain the reproductive processes in *T. hemprichii* (Pedersen et al., 2016).

Anthropogenic pollution such as waste water discharge, land run-off and aquaculture expansion can cause nutrient enrichment, eutrophication, turbidity and resuspension of sediments in the water column (Sachithanandam et al., 2020). These anthropogenic inputs can also increase the concentration of toxic trace elements like Pb and Cd in the coastal water and sediments of ANI Islands that can have negative impacts on the *T. hemprichii* population (Nobi et al., 2010; Mishra and Kumar, 2020). This nutrient enrichment and its negative effects on *T. hemprichii* population will mostly depend on the sediment grain size fractions, as finer sediment grain size (<65μm) will help in retaining these nutrients and heavy metals (Mishra and Kumar, 2020). Interestingly, the disturbed site of Burmanallah had higher fine grain particles than the undisturbed site (Mishra and Kumar, 2020; Mishra and Apte, 2020), that may have influenced the leaf growth rates at this site leading to lower leaf lengths among the three study locations (McDonald et al., 2016; Kelly and Dunton, 2017; Mishra and Apte, 2020). However, under these conditions *T. hemprichii* invests more in reproductive processes to migrate towards a more suitable environment through horizontal migration and seed dispersal (Lawrence and Gladish, 2019; Mishra and Apte, 2020). However, under these conditions *T. hemprichii* invests more in reproductive processes to migrate towards a more suitable environment through horizontal migration and seed dispersal (Lawrence and Gladish, 2019). Similar phenomenon has also been observed for *T. testudinum* from the coast of USA (McDonald et al., 2016; Kelly and Dunton, 2017).

Positive correlation of RE with plant size and growth suggests that seagrass sexual reproduction is a collective process of the plant, where the plant morphometric features such as horizontal rhizome diameter, elongation rate, leaf length and shoot biomass contribute to the reproductive process (Rollon et al., 2001; McDonald et al., 2016; Lawrence and Gladish, 2019). Positive correlation with rhizome diameter suggests that *T. hemprichii* having thicker rhizomes responded positively to disturbance and increased its RE (Cabaço and Santos, 2012). The rhizome diameter is closely related to plant size and shoot mass (Marbà and Duarte, 1998) as a result the positive response to disturbance through increased RE is related to quantity of energy stored in the rhizome and the plant size (Kuo and Den Hartog, 2006). Higher plant size will lead to more energy storage and will contribute to higher RE, that can result in higher flowering/fruit production during the disturbed conditions as flowering/fruit production are energy demanding process (Hirose et al., 2005; Karlsson and Méndez, 2005). Higher shoot biomass also helps the plant not only in storing energy, but also helps in regulating the nutrient content (C and N) in the meadow (Stapel et al., 1997) which eventually helps the plant in increasing its RE. This phenomenon has been observed for *T. hemprichii* from the coast of Indonesia (Stapel et al., 1997). Shoot density of seagrasses also play an important role in retaining the seagrass fruits (Meysick et al. 2019) and the higher reproductive density (than the undisturbed sites) at the disturbed sites (site 1) can be a result of successful retention of *T. hemprichii* fruits even with low shoot density. Similar cases of increase in RE resulting in higher fruit production had been observed for *T. hemprichii* population along the coast of Southern Thailand (Tongkok et al. 2017) and the Indonesia (McMahon et al. 2017) and for mixed meadows of *Halodule uninervis*, T*. hemprichii*, *Cymodocea rotundata* and *E. acoroides* of Philippines (Olesen et al. 2004) in Andaman Sea.

Though an increase in reproductive ecology (production of fruits/flowers) has been reported for T. *hemprichii* around the Andaman Sea (Agawin et al., 2001; Rollon et al., 2001; Olesen et al., 2014; McMahon et al., 2017), we report for the first time about the increase in RE of *T. hemprichii* under the influence of various habitat disturbances. *T. hemprichii* morphometric and physiological parameters have been used as an indicator to various disturbances (Roca et al., 2016; Ali et al., 2018; Zulfikar et al., 2020; Mishra and Apte, 2020) and we provide evidence that *T. hemprichii* RE can be used as an indicator of coastal disturbances around the Andaman Sea. *T. hemprichii* reproductive ecology varies spatially (Rollon et al., 2001; McDonald et al., 2016) and we observed the RE of *T. hemprichii* being site-specific and varying between the locations of ANI. This variation in RE are dependent on the site-specific environmental parameters, such as temperature, nutrients, light conditions and wave exposure (Agawin et al., 2001; Ali et al., 2018) and the population structure (Mishra and Apte, 2020) of the plant during the disturbances that eventually decide the response of the plant.

Our results will contribute strongly to increase the knowledge on *T. hemprichii* RE around the Andaman Sea and the Indian Ocean bioregion. This information will also add new data on *T. hemprichii* reproductive ecology from the coast of India (Table 2). However, a collective study around the Andaman Sea bioregion is necessary to generate a clear picture of the response and adaptation of this keystone species for better management and conservation measures, as T. hemprichii population is under decline at certain locations of ANI (Mishra and Apte, 2020).

## 5. Conclusion

We report for the first time about the reproductive ecology and RE of *T. hemprichii* from the coast of India. We observed a positive response of seagrass to habitat disturbance through increase in RE. Our results provide evidence that *T. hemprichii* can adapt to habitat disturbance through plasticity in RE, even though this response can be site-specific. This plasticity in adaption to habitat disturbances through an increase in RE can enhance the plants ability to withstand unfavourable conditions, improve seedling dispersal and settlement mechanisms towards favourable habitats and consequently increase the genetic diversity of the population (Hughes and Stachowicz, 2004; Coyer et al., 2004; Ehlers et al., 2008; Hughes et al., 2009; McDonald et al., 2016). Increase in genetic diversity of the plant will also contribute to maintain the various ecological functions and ecosystem services seagrasses provide (Ehlers et al., 2008; Hughes and Stachowicz, 2009; McKenzie et al., 2020). Our findings indicate that the RE of *T. hemprichii* can be used as a useful indicator of habitat disturbances (Table 2) along with various other morphological and physiological parameters previously used (Agawin et al., 2001; Ali et al., 2018). Our results suggest a 4-fold increase in RE to habitat disturbances can be used as a metric/ ecological indicator, which is also recommended for various other seagrass species (Cabaço and Santos, 2012). We provide initial information about the seagrass meadows of ANI that are under stress due to various habitat disturbances and call for proper mitigation measures from the local environmental managers to prevent seagrass loss. We propose more studies around the coast of India to use RE as a tool to assess seagrass ecosystems, as loss of these valuable ecosystems will lead to loss of 24 different types of ecosystem services (Unsworth et al., 2018) that benefits both seagrasses associated biodiversity and the livelihood of coastal human communities.

## Acknowledgement

We are grateful to the PCCF, wildlife for providing the required official help during the sampling programme. We are thankful to P M Ishaaq and Sumantha Narayana for the help during field work. We are thankful to Meena R Maithreyi for her support in making the manuscript better.

## Tables

Table 1. Mean ± S.D (standard deviation) of environmental variables of seawater and sediment grain size fractions of disturbed (Site1) and undisturbed (Site2) sites of Neil, Havelock Islands and Burmanallah locations of ANI. Water temperature (T°C).

Table 2. Effects of various disturbances (natural and anthropogenic) on the seagrass reproductive effort of *Thalassia testudinum* and *Thalassia hemprichii* species from around the world and around bioregion surrounding Andaman Sea. Reports that mentioned only the reproductive ecology without any disturbances/stressors were excluded.

## Notes

### Competing Interest Statement

The authors have declared no competing interest.

